# Tracking long-term natural changes in hippocampal excitatory to inhibitory synapses from spikes

**DOI:** 10.64898/2026.05.27.728158

**Authors:** Naixin Ren, Abhijith Mankili, Ian H. Stevenson

## Abstract

Although long-term synaptic plasticity is often studied using controlled electrical or optogenetic stimulation, it also occurs spontaneously during natural, ongoing brain activity. Here, using large-scale spike recordings in mice from the Allen Institute Visual Coding Neuropixels dataset, we examine to what extent fluctuations in putative synaptic efficacy can be tracked and explained by models of activity-dependent long-term plasticity. We first detect putative excitatory synaptic connections within the hippocampus based on cross-correlations between the spike trains of thousands of pairs of neurons. Most of these putative connections are between presynaptic neurons with broad spike waveforms and postsynaptic neurons with narrow spike waveforms and are consistent with synapses from excitatory to inhibitory units. For the subset of pairs where a transient, excitatory effect was detected, we use a model-based approach to track fluctuations in synaptic efficacy. Previous work found that these fluctuations can be partially predicted from pre- and postsynaptic firing rates and models of short-term plasticity. Here we model naturally occurring long-term potentiation and depression using rate-based learning rules. We find that modeling the covariance of pre- and postsynaptic activity improves prediction of efficacy fluctuations. We also examine synaptic changes associated with hippocampal sharp-wave ripples, but do not find clear evidence of systematic SWR-associated changes for the putative synapses studied here.

## Introduction

Long-term potentiation and depression (LTP and LTD) are fundamental mechanisms underlying learning and memory, and most neural systems exhibit activity-dependent changes in synaptic strength following pre- and postsynaptic spiking lasting minutes to hours (Malenka and Bear, 2004). Traditional computational models of long-term plasticity (Hebb, 1949; Abbott and Nelson, 2000) have emphasized two broad types of changes: Hebbian plasticity, where correlated pre- and postsynaptic firing leads to increased synaptic strength, and anti-Hebbian plasticity, where correlated firing reduces synaptic strength. Using controlled stimulation protocols *in vitro* and *in vivo*, the detailed molecular, physiological (Malenka, 1994), and computational (Feldman, 2020) underpinnings of long-term plasticity have been well characterized in many systems. One expectation from this work is that long-term fluctuations in synaptic strength should also occur during free behavior in response to uncontrolled, ongoing neural activity (Dobrunz and Stevens, 1999). However, modeling ongoing fluctuations in synaptic strength is a challenge, due to variability not just in measurements of post-synaptic strength but also the variability in the input and the presence of potential unobserved confounding variables. During free behavior, neuromodulation, complex activity patterns of multiple presynaptic inputs, and changes in brain and behavioral states (Bramham and Srebro, 1989) may also alter the context of synaptic changes, potentially engaging different plasticity mechanisms or modulating their expression. Here we develop data analytic approaches for tracking long-term fluctuations in synaptic strength in vivo during natural, ongoing brain activity, using large-scale spike recordings.

Many previous studies have outlined how synapses can be characterized from simultaneous pre- and postsynaptic spiking (Perkel et al., 1967; Swadlow and Lukatela, 1996), and recent work has applied these methods to large-scale spike recordings (Barthó et al., 2004; Stevenson, 2023; Kobayashi and Shinomoto, 2025). Rather than estimating synaptic strength from post-synaptic potentials or currents as in intracellular recordings, here we detect putative pairs of synaptically connected neurons from spike recordings based on cross-correlations and estimate the change in excess postsynaptic spike probability, the efficacy, in the short window following presynaptic spikes. In previous work, we found that the efficacies of putative synapses in behaving animals show slow natural fluctuations over time that are correlated with the time-varying presynaptic firing rate (Ren et al., 2022) with positive correlations for some neuron pairs and negative correlations for others. Although these effects occur over a timescale of minutes, in most cases they are consistent with short-term plasticity (STP) where short-term depression and facilitation lead to predictable changes in steady-state efficacy as presynaptic rates fluctuate. Here we investigate the role of long-term plasticity mechanisms and determine to what extent rate-dependent models of long-term potentiation and depression (LTP and LTD) can account for additional unexplained variance in observed synaptic efficacy fluctuations.

Here we focus on hippocampal excitatory-to-inhibitory (E-I) synapses in mice. Activity and plasticity at these synapses has been shown to be essential for regulating neuronal excitability and hippocampal network dynamics (Klausberger and Somogyi, 2008; David and Topolnik, 2017) as well as memory (Topolnik and Tamboli, 2022). Many previous studies have examined long-term plasticity at these synapses using intracellular recordings, and hippocampal E-I connections show substantial diversity in plasticity (Kullmann and Lamsa, 2007; Pelletier and Lacaille, 2008 p.14). LTP at E-I synapses onto both parvalbumin-positive (PV) interneurons (Le Roux et al., 2013) and somatostatin-positive (SOM) interneurons (Honoré et al., 2021) can be induced in slice experiments by high-frequency stimulation. LTD can be also be induced by stimulation that typically induces LTP at E-E synapses (McMahon and Kauer, 1997; Laezza et al., 1999). Hebbian and anti-Hebbian forms of plasticity have both been observed, and the time course and parameters necessary for induction appear to be input specific, use different molecular mechanisms, and vary by cell type and location (Kullmann and Lamsa, 2007). However, in most cases, it is unclear to what extent predictions of LTP or LTD from *in vitro* studies generalize to explain ongoing fluctuations in synaptic strength *in vivo*. Here we use an observational approach and aim to track and model fluctuations in synaptic strength at E-I connections *in vivo* during natural, uncontrolled brain activity. Since structured pairing events are a key component of many LTD/LTP studies, we also examine the effects of hippocampal sharp-wave ripples (Buzsáki, 2015; Joo and Frank, 2018) where both pre- and post-synaptic neurons repeatedly fire together (Csicsvari et al., 1999; Noguchi et al., 2022).

To characterize the fluctuations in the efficacy of putative hippocampal E-I synapses from spikes we take two approaches: an empirical approach that focuses on estimating and tracking time-varying changes in synaptic weight, and a computational approach that aims to determine whether the fluctuations are consistent with parametric models of long-term plasticity. Although there are many computational models of long-term plasticity (Morrison et al., 2008), here we focus on the Bienenstock-Cooper-Munro (BCM) rule (Bienenstock et al., 1982). The BCM rule models long-term weight changes based on the activity and coactivity of the pre- and postsynaptic neurons, and allows both Hebbian and anti-Hebbian changes with both LTP and LTD. The BCM rule appears to qualitatively explain many features of hippocampal LTP and LTD (Hulme et al., 2012), and, since the rule depends exclusively on pre- and postsynaptic rates, we can directly evaluate to what extent covariation in rates is associated with observed efficacy fluctuations. During ongoing activity, the firing rates of both pre- and postsynaptic neurons vary over time and their coactivity is modulated by stimuli, behavior, and brain state (Salinas and Sejnowski, 2001; Averbeck et al., 2006; Churchland and Abbott, 2012). Here we aim to identify to what extent the fluctuating relationship between pre- and postsynaptic firing rates during ongoing brain activity explain observed efficacy fluctuations.

We first detect and characterize putative hippocampal E-I synapses using large-scale multi-electrode recordings. We then introduce methods for estimating slow fluctuations in efficacy over time and for estimating observed changes in efficacy before and after discrete events. We find that for individual, putative synaptic connections, slow fluctuations in synaptic efficacy appear that are consistent with both Hebbian and anti-Hebbian plasticity. Fitting rate-based models to the observed fluctuations, we find that efficacy changes are not explained by STP alone, and modeling the correlation of the pre- and postsynaptic spikes accounts for additional long-term changes in the synaptic weight. Finally, we examine how hippocampal sharp-wave ripples (SWRs) might influence LTP/LTD. We find that although the firing rate of both pre- and postsynaptic neurons increases during SWRs for almost all E-I pairs, there do not seem to be consistent alterations in efficacy during ongoing brain activity. Altogether, these results illustrate how short- and long-term plasticity may both explain natural fluctuations in synaptic efficacy in behaving animals.

## Results

Our aim is to identify putative hippocampal synapses from large-scale spike recordings, to track fluctuations in putative efficacy over time, and to determine to what extent activity-based models of long-term plasticity can account for these fluctuations. In controlled experiments (Fig 1A) the activity of the pre- and postsynaptic neurons are directly manipulated, and the effects of the manipulation can be measured and averaged to characterize rate-dependent LTP and LTD. During natural, ongoing brain activity, the same LTP/LTD mechanisms may be occurring (Fig 1B) but require a more intensive analysis to disentangle. Here we test whether the BCM rule (Fig 1C), with either Hebbian or anti-Hebbian learning rules, can account for fluctuations in efficacy that may occur spontaneously.

**Figure 1:**
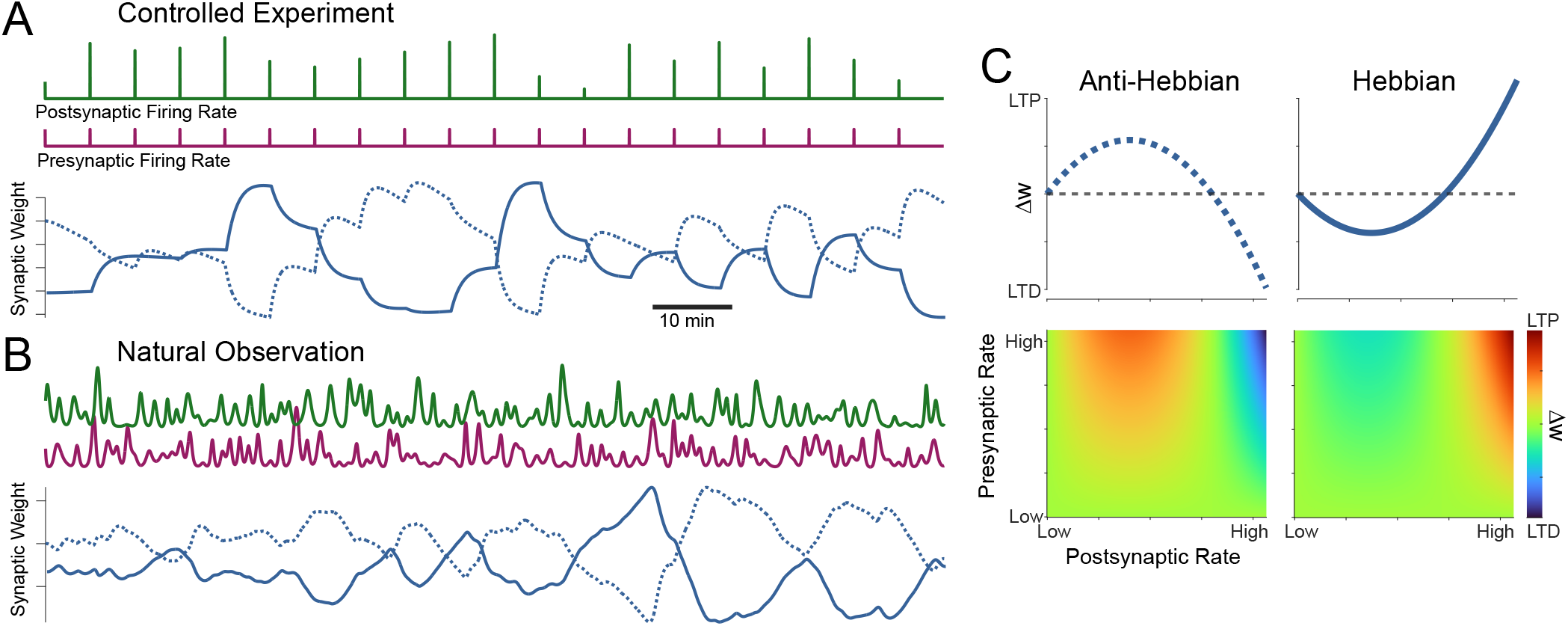
Studying rate-dependent long-term plasticity during natural observation. A) Controlled manipulation of pre- and postsynaptic rates has been shown to produce both long-term potentiation (LTP) and long-term depression (LTD). Pairing can result in Hebbian (solid) or anti-Hebbian patterns of LTP/LTD. B) Patterns of rate pairing may lead to spontaneous fluctuations in synaptic strength when pre- and postsynaptic firing rates are uncontrolled. C) Synaptic weight changes for a BCM model with anti-Hebbian (left) or Hebbian (right) parameter settings at a fixed presynaptic rate (top) and for all combinations of pre- and postsynaptic weights (bottom). These models were used to generate the synaptic weight traces in (A) and (B).

### Putative hippocampal synaptic connections

We focus here on putative hippocampal synapses identified from large-scale spike recordings in the Allen Institute – Visual coding Neuropixels dataset (see Methods). The pre- and postsynaptic neurons examined here are recorded from hippocampal areas CA1, CA3, dentate gyrus (DG), subiculum (SUB), and prosubiculum (ProS). Across 52 recordings, there are 6234 units from these brain areas with SNR>2 and more than 1000 spikes. The distribution of units and pairs is not uniform across the areas, and the majority of units come from CA1 (n=3811) and DG (n=870).

Using a two-stage synapse detection method we identify putative synapses with cross-correlograms (CCGs) that are consistent with the biophysics of excitatory synaptic transmission (Fig 2). Altogether, our screening method identifies n=666 pairs of putative excitatory synapses within and between these areas. Out of 870K possible pairs 6501 pairs passed an initial screening based on hypothesis testing (with Benjamini-Hochberg correction, false discovery rate 0.1), and 666 were identified as having sufficient goodness-of-fit and synaptic latencies and time-constants consistent with excitatory synaptic connections (see Methods). Of these putative excitatory synapses, 578 are intra-area connections and 486 are within CA1 (Fig 2A).

**Figure 2:**
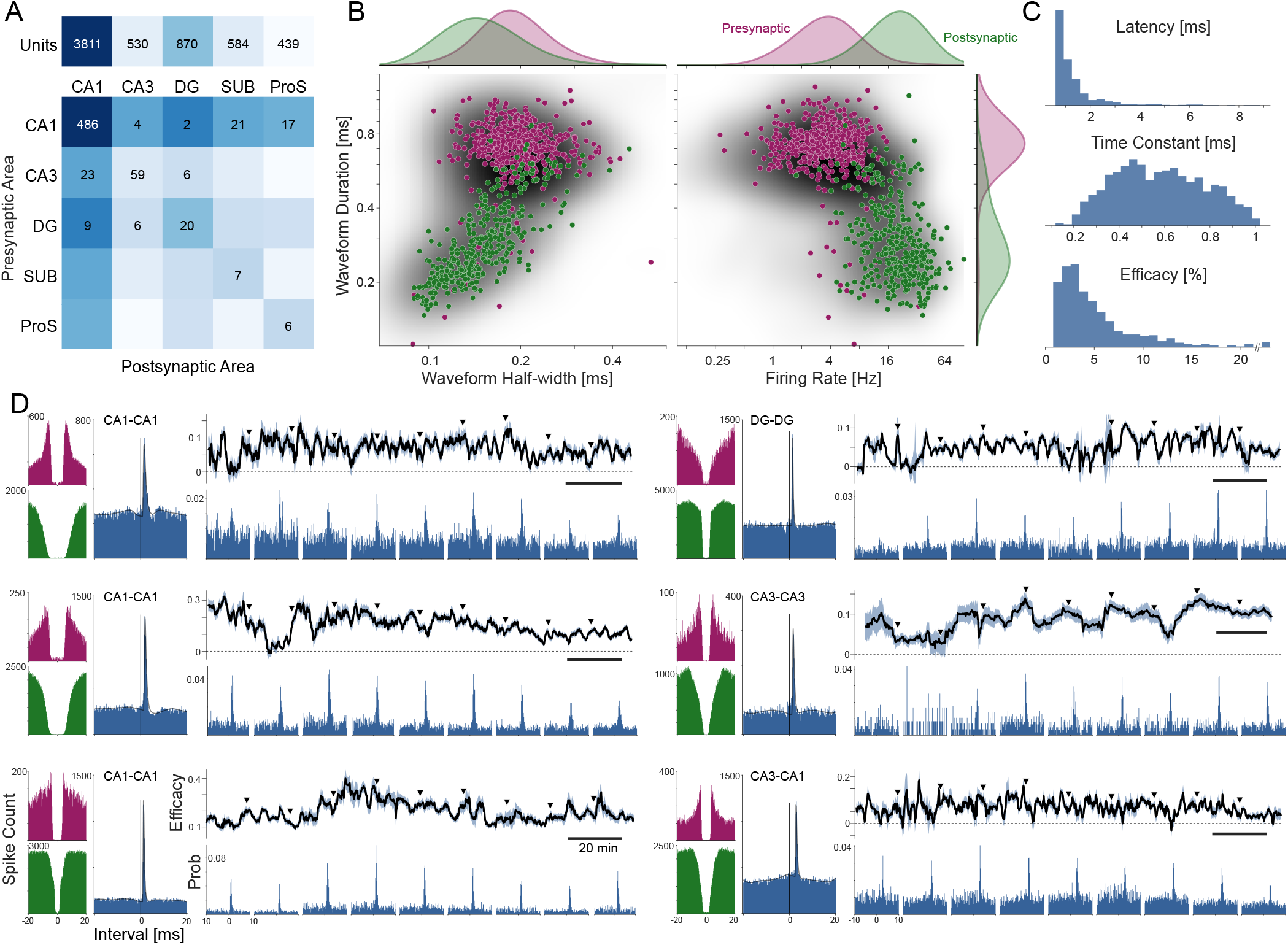
Putative hippocampal synapses and long-term fluctuations in efficacy. A) Counts of units and detected putative synapses by hippocampal area. Color in the connectivity matrix indicates the relative number of possible pairs between regions. B) Waveform statistics for the putative pre- (purple) and postsynaptic (green) neurons. Gray distribution in background denotes the statistics for all hippocampal units. C) Model-based estimates of synaptic latency, time constant, and efficacy for the putative synaptic connections. D) 6 examples of putative synapses (each from a different recording) and their fluctuating synaptic efficacy over the duration of the recording. The overall cross-correlogram (blue), as well as the overall, auto-correlograms for the pre- (purple) and postsynaptic (green) units are shown at left. The grey curve overlaid on the cross-correlogram denotes the model fit. Efficacy traces and errorbars denote estimates and standard errors from a model-based moving window method whose size is optimized by cross-validation. The cross-correlograms below indicate windowed data ±5 minutes around the time points indicated by triangles.

The majority of detected pairs represent putative synapses from units with a broad waveform to units with a narrow waveform. The median waveform duration for the putative presynaptic units is 0.73 ms, while the median duration for the postsynaptic units is 0.26 ms. As previous studies have noted, units with narrow waveforms also tend to have higher firing rates (Connors and Gutnick, 1990), and this pattern occurs here as well. Presynaptic units have median firing rate of 3.7 Hz (2.3-5.6 Hz, Q1-Q3), while the postsynaptic units have median firing rate of 20.3 Hz (13.1-29.3 Hz). These results are consistent with E-I connections where excitatory units project to inhibitory units (Connors and Gutnick, 1990; Barthó et al., 2004; Trainito et al., 2019).

We fit a convolutional model to the overall cross-correlograms to characterize the parameters of the putative synaptic effects (Fig 2C). This model (see Methods) directly fits the correlogram with a fast, transient component (alpha function) that captures a synaptic effect and a slower component (B-spline basis) that captures the shape of the baseline postsynaptic activity around each presynaptic spike. Here the median latency is 0.94 ms (0.86-1.37 ms, Q1-Q3) with a small set of pairs (27 of 666) having latency >4 ms, median time constant is 0.57 ms (0.42-0.73 ms), and median efficacy is 3.7% (2.4-6.5%) with a small set of pairs (12 of 666) having estimated efficacy >20%.

To study how the strength of putative synapses changes over time we split the recording into smaller time periods. We find that, while the cross-correlograms calculated over short segments of data often have peaks with the same latency and time constant as the overall correlogram, there are fluctuations in the apparent efficacy over time (Fig 2D). To track fluctuations, we first fit the latency and time constant for the overall correlogram and then use a moving window to estimate local efficacy by updating the weight of the fast model component. We optimize the window size (between 1-10 minutes) for each putative synapse by maximizing the cross-validated log-likelihood of the moving CCGs (5-fold cross-validation with 10s test periods). This approach allows the window size to vary based on the precision of efficacy estimates for each pair. Units with lower firing rates require longer window sizes for accurate estimates, but, for each pair, the optimal window size avoids over-smoothing (missing observable fluctuations) as well as under-smoothing (tracking noise). The optimal window size has median 126 s (45-246s, Q1-Q3), and the median coefficient of variation for the estimated efficacy fluctuations is 0.83 (0.57-1.50). We observe consistent fluctuations in efficacy for all types of inter- and intra-region putative synapses detected here (Fig 2D).

Estimates of time-varying efficacy do have limited precision. The number of postsynaptic spikes that contribute to the efficacy estimate is typically a small fraction of the total postsynaptic spikes. The contribution (the proportion of postsynaptic spikes that fall within the fast, transient effect and are estimated to be above the baseline) is much smaller than the efficacy (median 0.7%, 0.4-1.3% Q1-Q3) for the pairs analyzed here. This amounts to only 0.14 “excess” spikes per second for the typical (median) pair. To evaluate the reliability of these efficacy fluctuations we thus consider a control based on altering or shuffling the postsynaptic spikes that contribute to the efficacy estimate. In particular, we identify the subset of postsynaptic spikes that occur within the time period defined by the fast component of our model. For each of these postsynaptic spikes we identify its interval relative to the most recent presynaptic spike. We then reassign these postsynaptic spikes to occur with the same interval after a randomly selected presynaptic spike from the whole recording. This generates a type of negative control where the presynaptic spikes and overall correlogram do not change, and the postsynaptic spikes are mostly unchanged except for the small fraction that occur within the time period of putative synaptic transmission. However, after shuffling, any fluctuations in the efficacy over time are due to chance, since each presynaptic spike has the same probability of being followed by excess postsynaptic spikes (Fig 3A).

**Figure 3:**
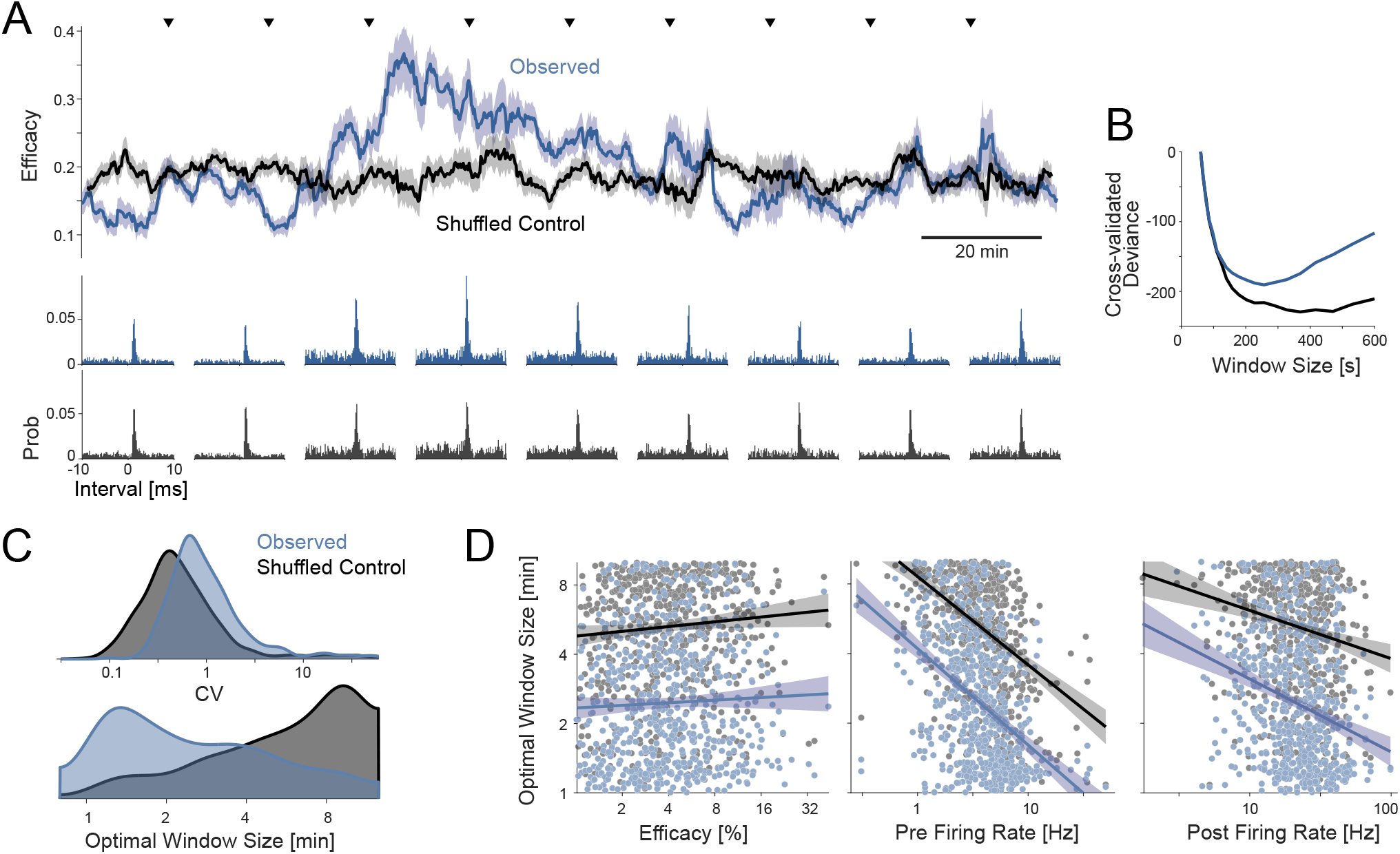
Efficacy fluctuations estimated from shuffled spikes. A) Example correlograms and efficacy traces from observed data (blue) and a shuffled control (gray). Triangles denote measurement times for cross-correlograms shown below. Errorbars denote estimates and standard errors from a model-based moving window method. This specific example corresponds to the CA1-CA1 pair in Fig 2D, bottom. B) Window size is optimized for each pair by minimizing the local cross-validated deviance. The result for the example in (A) is shown here for observed (blue) and shuffled (gray) spikes. C) Distribution of coefficient of variations (CV) for observed efficacy fluctuations and shuffled controls for all neuron pairs. D) Optimal window size is associated with the efficacy and rates of the pre- and post-synaptic neurons. Dots denote individual putative synapses. Lines denote linear best fit to the log-log plot for the observed (blue) and shuffled (gray) putative synapses, respectively, with 95% confidence bands.

Optimizing the window size for pairs that have been shuffled with this control results in estimated efficacy fluctuations that are more stable with longer optimal windows sizes (median 370s, 228-600s Q1-Q3) compared to the observed, unaltered data (Fig 3C). The efficacy fluctuations estimated from the unaltered data have a higher coefficient of variation compared to the negative controls (median 0.47, 0.3-0.86) at their respective optimal window sizes (Fig 3C). As expected, the window size also depends on the pre- and postsynaptic rates, with lower rates leading to longer optimal window sizes (Fig 3D). One major factor affecting the estimates of time-varying efficacy is the time-varying baseline. When the postsynaptic rates vary over the course of the recording this can alter the efficacy estimates, and these baseline fluctuations may be one reason why the estimated efficacy fluctuations for the controls still show some variation. However, these controls provide evidence that efficacy fluctuations observed in the original data are larger than expected by chance.

### Observed relationships between pairing and long-term efficacy changes during ongoing activity

In controlled experiments, it is possible to directly observe the causal impact that specific combinations of pre- and postsynaptic firing rate have by stimulating and subsequently measuring the evolution of the synaptic strength. For natural rate fluctuations, however, estimating the causal impact of combinations of firing rates requires disentangling the confounding effects of many other uncontrolled factors, such as short-term plasticity and changes in excitability. Combinations of pre- and postsynaptic rates occur continuously with an uncontrolled distribution and with strong temporal correlations. As an initial analysis, we nonetheless measure the associations between pre- and postsynaptic rates and subsequent efficacy changes using a time-weighted cross-correlogram.

We first estimate the firing rates for each neuron in 100ms bins and smoothed using an optimized kernel density estimate (Shimazaki and Shinomoto, 2010). We then identify points in time where the firing rates of the putative presynaptic neuron and postsynaptic neuron occur in specific combinations (e.g. when both are high or both are low). Using these time points we weigh the spikes before and after these events and construct a modified correlogram (see Methods). Here weighting functions are generated by filtering the event times using an alpha function (τ_*r*_ = 5 min) – filtering forward for “after” and backward for “before. The correlograms using weighted-before and weighted-after spikes typically have the same latency and time constant but sometimes differ in their efficacies. The difference between the efficacies before and after provides an estimate of Δeff for that specific pre-post firing rate combination (Fig 4). Evaluating the weighted-before and weighted-after correlograms for many combinations of pre- and post-synaptic rate we can visualize how these events were associated with weight changes and estimate a map Δ*eff*(*r*_*pre*_, *r*_*post*_) describing observed firing rate-based efficacy changes. To standardize across pairs, we estimate this map using the ranks/percentiles of the pre- and postsynaptic firing rates.

**Figure 4:**
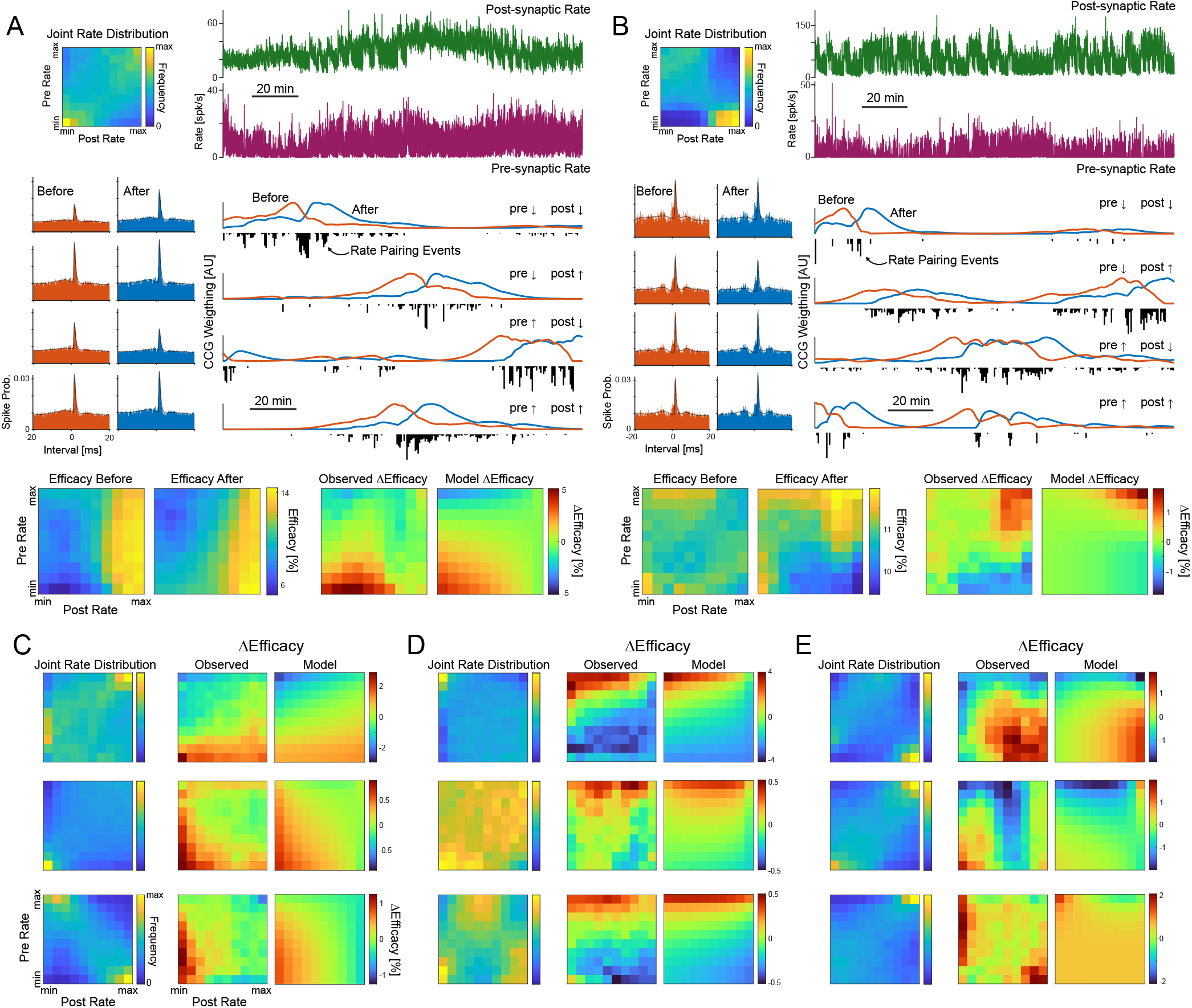
Observed changes in efficacy following natural rate pairing events. A and B show example analyses for two putative pre- and post-synaptic neuron pairs within CA1. Both pre- and post-synaptic rates fluctuate over the course of the recording (top right) and rate combinations occur with different frequencies (top left). For each rate combination (e.g. pre low/post low or pre high/post high) we identify the time of these “rate pairing events”. We then generate two CCG weighting functions to describe time points “before” and “after” pairing events (bottom right) and use these to estimate CCGs (bottom left) and efficacy before and after each particular combination of rates. In some cases, the observed changes in efficacy are consistent with anti-Hebbian (A) or Hebbian (B) plasticity, and we model these effects by fitting the observed efficacy changes with a BCM rule. C-E) Additional examples of anti-Hebbian-like (C) and Hebbian-like (D) plasticity, as well as, examples that are not well-described by the BCM. All examples here have high split-half reliability (*r*>0.5). Top examples in C-D are CA3-DG all others are pairs within CA1.

We find that the efficacy changes following naturally occurring combinations of high pre- and post-firing rates can be negative, consistent with LTD and anti-Hebbian plasticity (Fig 4A, C), or positive, consistent with LTP and Hebbian plasticity (Fig 4B, D). In many cases, the same pair will have positive Δ*eff* for some rate combinations and negative for others. The maps are diverse (Fig 4C-E) and often nonlinear functions (Fig 4E). Here we fit three potential learning rules: 1) a non-Hebbian rule, linear in the pre- and postsynaptic rates, 2) a covariance rule, which adds a second-order term, and 3) a BCM rule. We find that the non-Hebbian, linear rule explains 35±23% of the variance (mean±sd) in the efficacy change maps, while a covariance rule explains 41±23% and a BCM rule explains 45±23%.

The recordings here are ~2hrs on average and one possibility is that this is not sufficiently long to reliably observe LTP and LTD. To evaluate the reliability of the efficacy change maps we use split-half validation, comparing the map for the first half of the recording to the map for the second. The examples in Fig 4 represent cases with high split-half reliability (*r* =0.48 [0.31,0.61] for Fig 3A, *r* =0.70 [0.58,0.79] for Fig 3B, 95% CI, and *r* > 0.5 for Fig 3C-E). Many pairs (n=63 of 666) have *r*>0.5 but the correlation between the first and second half estimates across all pairs is low on average 0.02±0.36 (mean±sd). This may be due to the recording length as well as the fact that these maps do not address potential confounds and do not model the accumulation of LTP/LTD over time. We, thus, turn to alternative models that may be able to more accurately account for *eff*(t).

### Linear models of fluctuating efficacy

In previous work, we showed how the fluctuations in pre- and postsynaptic firing rates, by themselves, can account for some long-term fluctuations in efficacy. In particular, short-term depression and facilitation can directly induce negative or positive correlations between the presynaptic firing rate and the efficacy on timescales of seconds to minutes (Ren et al., 2022). Here we use a linear model of these first-order rate-related effects and then add higher-order effects that can account for potential changes due pre-post pairing.

Using the optimal window size estimates for putative synaptic efficacy, we generate predictor variables on the same timescale. For the rate-model we use the log pre- and postsynaptic firing rates estimated with optimal bandwidth (Shimazaki and Shinomoto, 2010) then smoothed to match the window size for efficacy estimates. To model rate-based LTP/LTD we then add additional higher-order terms that are filtered to mimic the accumulation of long-term plasticity over time: an accumulated covariance term *r*_*pre*_, *r*_*post*_ * *w* and an accumulated 3^rd^-order term 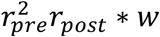, where *w* denotes an alpha function with time constant τ and * denotes convolution. Using these predictors we then fit the observed efficacy using linear regression and weight each time point by the uncertainty of the efficacy estimate.

We find that the Rate+BCM linear model can, in many cases, capture efficacy fluctuations more accurately than the first-order rate model alone (Fig 5 A, B). Using the coefficients for the BCM effects – the terms modeling 2^nd^ and 3^rd^ order rate effects – we can also examine the rate-based learning rule for this model Δ*w*(*r*_*pre*_, *r*_*post*_). Although these rules, by design, match the BCM, we find that the linear model with accumulation typically does not align well with the observed Δ*eff* estimated by the weighted correlograms (Fig 5 C, D). However, the coefficients of the model tend to follow a consistent pattern across pairs and regions (Fig 5E). The 1^st^ order pre- and postsynaptic log-rate coefficients tend to be positive, while the 2^nd^ order accumulated covariance coefficients tend to be negative. There is substantial variability in the coefficients across pairs. However, under this model, most pairs would be described as anti-Hebbian. We do find that the results are fairly robust to the timescale on which the BCM effects accumulate (Fig 5F). Adding the 2^nd^ and 3^rd^ order terms improves the average variance explained for all 10s < τ <360s, and optimizing the time constant for each pair separately gives the greatest accuracy (27±2% variance explained for the Rate+BCM median±SE, compared to 16±1% for the 1^st^ order model). Due to overfitting, adding covariates is expected to improve accuracy. However, we find that the AIC is lowest for the Rate+BCM model in 82% of pairs and lowest for the Rate+Cov model for 10% of pairs, suggesting that the accumulated higher-order covariates do contain predictive value.

**Figure 5:**
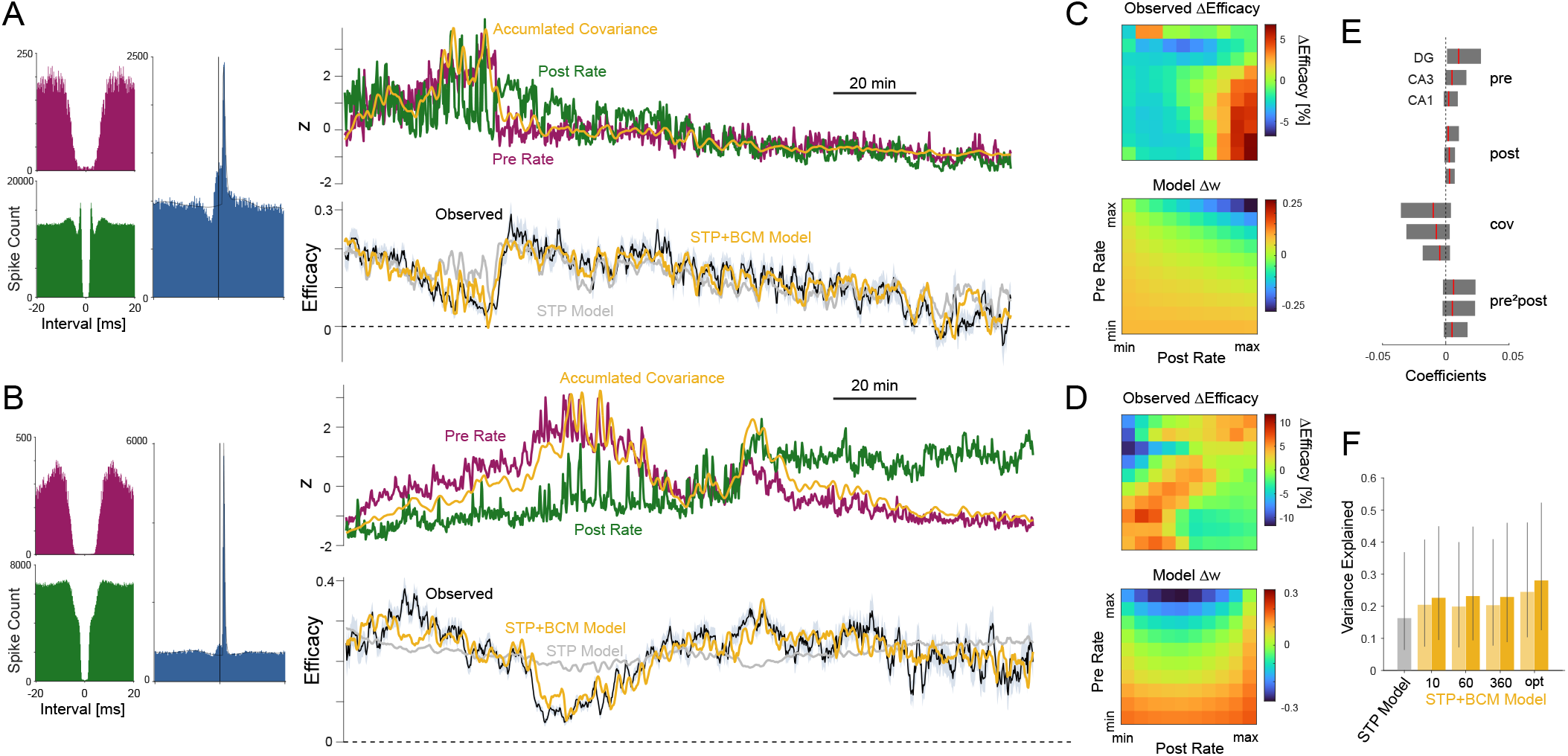
Dynamic linear models. A and B show two example putative synapses within CA1 with auto- and cross-correlograms (left), and time-varying rates and efficacy (right). Efficacy changes are correlated with and predicted by the changing firing rates of the pre- and postsynaptic neuron. Here we fit linear models with 1^st^-order rate terms (STP model) and, additionally, fit models with higher-order terms from the BCM model. Including the accumulated covariance often improves the fits. However, we find that the observed efficacy changes following pairing events typically does not match the apparent BCM effects (C and D). The coefficients for these linear models are largely similar across hippocampal areas (E). Boxes denote inter-quartile range, and median (red) across pairs conditioned on the location of the presynaptic unit. F) The STP+BCM linear model tends to out-perform the STP model in terms of variance explained. Light-yellow bars denote the covariance model. Average performance is not substantially affected by the accumulation time constant (10 vs 60 vs 360 s). However, fits are improved when the time constant is optimized for each pair (“opt”). Bars denote median, error bars denote inter-quartile range. Examples in A and B use a time constant of 30s.

### Sharp-wave ripples and fluctuations in synaptic efficacy

To examine the potential role of hippocampal sharp-wave ripples (SWRs) in the fluctuations of the synaptic efficacy we detect SWRs from the simultaneously recorded local field potentials (LFPs; see Methods). For each putative synapse, we detect ripples on the LFP channel closest to the pre-synaptic unit (Fig 6A) and analyze the firing activity of the pre- and postsynaptic neurons during the SWRs. Here the average number of SWRs detected for any given pair of units is 1141 ± 385 (mean ± SD), 6.9 ± 2.3 ripples/min with an average ripple duration of 26 ± 11 ms. We then examine the pre- and postsynaptic spiking relative to the onset of SWRs. As in previous studies, we find here that SWRs reflect moments of pre- and post-synaptic coactivation (Fig 6B-C). Putative presynaptic units on average increase their firing rate to 17.5 Hz during SWRs from 4.6 Hz during non-SWR times, while putative postsynaptic units on average increase their firing rate to 85.8 Hz during SWRs from 23.7 Hz during non-SWR times. These events could, thus, have the potential to generate alterations in efficacy via Hebbian or anti-Hebbian effects.

**Figure 6:**
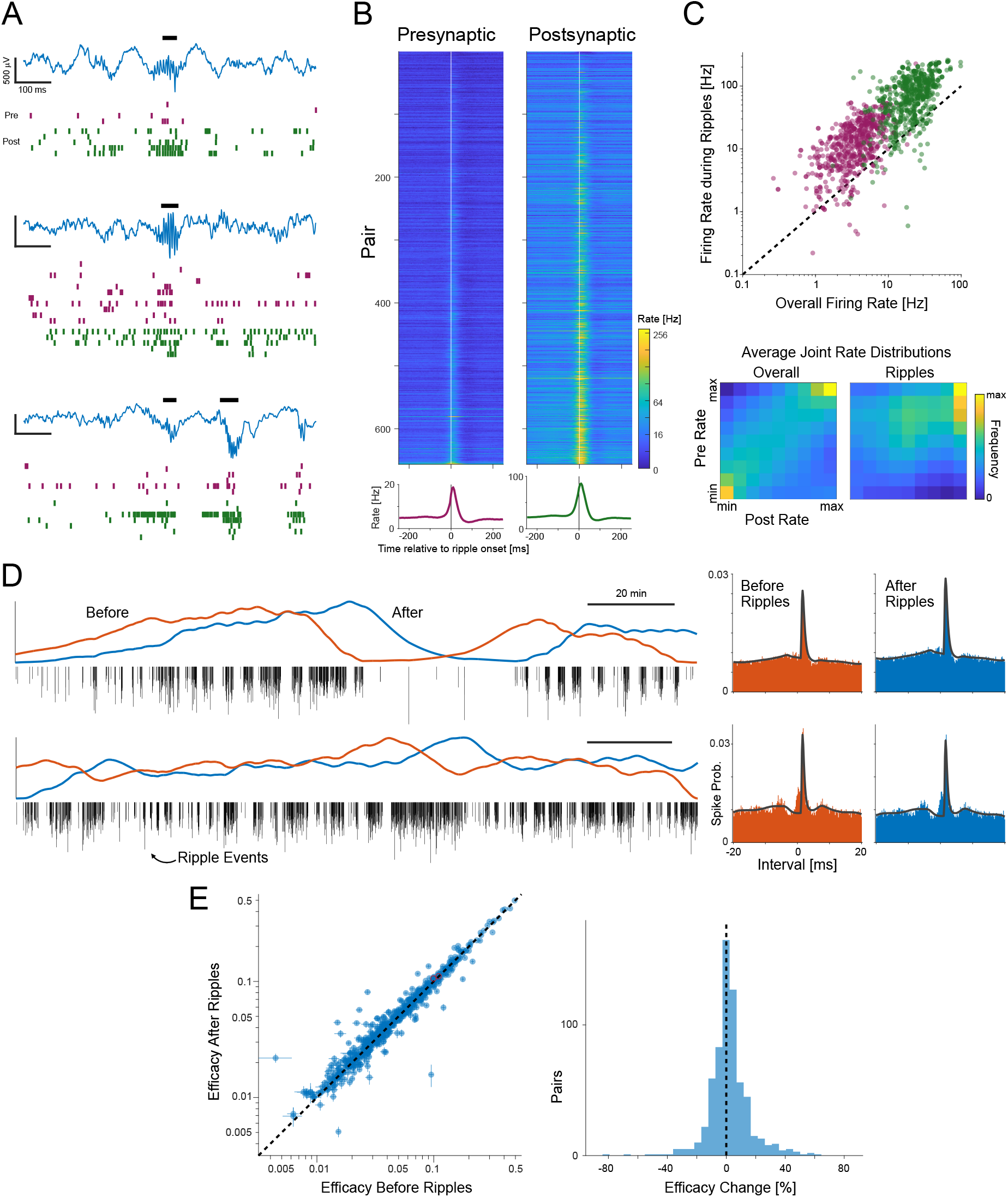
Pre-post pairing during sharp-wave ripples. A) Example traces of hippocampal local field potential activity (blue) where high frequency oscillations were detected (top) and the simultaneous spiking activity of hippocampal units recorded on the same probe and involved in putative E-I synapses. Examples are from three unique animals. Horizontal bars denote periods of time where the LFP met ripple criteria (see Methods). Since units can be involved in multiple pairs, the numbers of pre- (purple) and postsynaptic (green) units are not perfectly aligned. B) Average ripple-aligned firing rates for all pre- (left) and postsynaptic (right) units. C) Overall firing rates and firing rates during ripples for these units, and the corresponding pre-post joint rate distributions (rank transformed). D) Example ripple weighting functions for two recordings. Histogram denotes the timing of ripple events over the course of the recording. Blue and red traces denote forward (after-ripple) and backward (before-ripple) filtering of the ripple events. These functions are then used to weight spike counts for correlogram calculations. Two example putative synapses are shown at right. E) Observed efficacy changes across all pairs before and after ripples, estimated using the weighted CCGs.

Applying the same, weighted CCG method used above to describe efficacy change maps for all combinations of pre- and postsynaptic rates, here we assess changes before and after ripples. We generate weighting functions by filtering forward or backward with an alpha function (τ_*r*_ = 5*min*) and use a model-based approach to estimate the efficacy after and before the ripple events (Fig 6D). Although a subset of individual pairs show statistically significant changes (both increases and decreases) following SWRs (Fig 6E), on average there do not appear to be large, consistent changes. Across all pairs the efficacy increases slightly following SWRs by 1.3 ± 0.4% (median ± SE). Looking per-pair, we do find weak correlations between the SWR-associated efficacy change and the presynaptic rate change (*r* = −0.06) or postsynaptic rate change (*r* = −0.04) during the ripples and also a weak correlation between the before SWR efficacy and SWR-associated efficacy change (*r* = 0.06). These results suggest that some of the variability in SWR-associated efficacy changes may be due to variability in initial strength of putative synapses and variability in the extent of pairing that actually occurs during the ripples. However, the overall picture of SWR-associated efficacy changes seems to reflect only limited systematic efficacy changes. Instead, SWR-associated changes are highly variable, consistent with diverse efficacy change maps and a mixture of Hebbian and anti-Hebbian trends.

## Discussion

In this study we examined to what extent the natural fluctuations in synaptic efficacy of putative hippocampal excitatory connections can be predicted based on the relationship between the pre- and postsynaptic firing using large-scale extracellular spike recordings from awake, behaving mice. We identified 666 putative synapses with short latency, transient increases in postsynaptic spike probability following presynaptic spikes. The spike waveforms and rates of these pairs are consistent with hippocampal E-I connections, and we found that these connections often show substantial fluctuations in efficacy over the course of the recordings in the Allen Institute – Visual Coding Neuropixels dataset. We developed data analytic methods for estimating continuous changes in efficacy before and after environmental events using weighted cross-correlograms. We found that efficacy change maps for these connections can reliably show both Hebbian and anti-Hebbian effects. We then fit a dynamic model of time-varying efficacy and found that the higher-order effects in the BCM, including a term for the accumulated covariance of the pre- and postsynaptic rates, explain additional variability in efficacy fluctuations that is not explained by the individual firing rates alone. Finally, we estimated efficacy changes before and after sharp-wave ripple events. Here we did not find substantial, consistent changes. However, our results, altogether, suggest that activity-dependent LTP/LTD may shape efficacy fluctuations that are observed during ongoing brain activity.

Previous studies have shown that long-term plasticity can be induced in awake animals by repetitively pairing postsynaptic and presynaptic activity with electrical stimulation (Errington et al., 1997; Orr et al., 2001; Jackson et al., 2006; Tsanov and Manahan-Vaughan, 2009; Cooke and Bear, 2010) or with sensory stimuli (Meliza and Dan, 2006; Mu and Poo, 2006; Vislay-Meltzer et al., 2006). Connectivity also changes in the intact brain with optogenetic stimulation (McKenzie et al., 2021; Spivak et al., 2024) or electrical microstimulation (Rebesco et al., 2010). Natural long-term plasticity-like changes in synapses have also been discovered in behaving animals after repeated presentation of specific visual stimulus (Frenkel et al., 2006; Sale et al., 2011) and behavioral training (Gruart et al., 2006; Whitlock et al., 2006; Fedulov et al., 2007). These findings together suggest that long-term plasticity occurs naturally in behaving animals and has the potential to contribute to the observed, uncontrolled fluctuations in synaptic efficacy. Here we find that the ongoing fluctuations at hippocampal E-I synapses and can be partially described by a combination of rate-dependent effects, consistent with short-term plasticity, and coactivity-dependent effects consistent with a BCM-like learning rule.

Detecting and modeling putative synapses from spikes is a statistical challenge (Stevenson, 2023; Kobayashi and Shinomoto, 2025). Although here we rely on circumstantial evidence, such as waveforms and firing rates, to conclude that these putative synapses correspond to hippocampal E-I connections, our population of pairs is largely consistent with previous studies (Csicsvari et al., 1998, 1999; Marshall et al., 2002), including studies that have experimentally verified cell types and monosynaptic connectivity (English et al., 2017; Spivak et al., 2024). Previous work has also demonstrated how these synapses might fluctuate over time (McKenzie et al., 2021; Ren et al., 2022). However, one challenge in modeling plasticity from observational data alone is that it is difficult to average over or otherwise account for possible unobserved, confounding influences on synaptic strength. The fact that we only model a single putative presynaptic input among many may be one explanation for why the variance explained is relatively low. Modeling additional presynaptic inputs would likely improve postsynaptic spike prediction (Volgushev et al., 2015), but may also be necessary to account for heterosynaptic plasticity (Chistiakova et al., 2014) and the potential impact that multiple inputs (Spivak et al., 2024) have on driving efficacy fluctuations.

In addition to focusing on single neuron pairs, here we have also limited our analysis to focus on rate-dependent and coactivity-dependent fluctuations described by the BCM rule. Although this rule captures the basic phenomenology of Hebbian and anti-Hebbian plasticity and incorporates both LTP and LTD, there are many other types of long-term plasticity that have been described (Abbott and Nelson, 2000). Spike-timing-dependent plasticity (STDP) and the precise relative timing of pre- and postsynaptic spikes (Dan and Poo, 2004; Markram et al., 2012), likely play an important role in accounting for complex hippocampal synaptic dynamics (Debanne et al., 1997; Froemke et al., 2010). While under certain assumptions the BCM rule can approximate STDP (Izhikevich and Desai, 2003), recent results also suggest accounting for behavioral-timescale plasticity (Bittner et al., 2017) and broader calcium-dependence (Graupner and Brunel, 2012; Chindemi et al., 2022) may be necessary to accurately describe LTP/LTD patterns. While methods for estimating more flexible learning rules have been developed for both intracellular observations (Chindemi et al., 2022; Chen et al., 2023) and spikes (Stevenson and Kording, 2011; Linderman et al., 2014; Song et al., 2015; Robinson et al., 2016; Hem et al., 2023), validating and selecting between these models is still a challenge. One consideration that we have aimed to address here is the potential interaction between multiple plasticity mechanisms on multiple timescales (Costa et al., 2015; Wei and Stevenson, 2021). By accounting for both short- and long-term plasticity effects we are better able to explain efficacy fluctuations than with short-term, rate-dependent effects alone. However, for the models and ~2hr recordings used here, there is still substantial unexplained variability in efficacy fluctuations.

Robust models of long-term plasticity should also be able to account for changes in efficacy across and due to different brain states. Here we examined the potential role of sharp-wave ripples (SWRs) in efficacy fluctuations of putative hippocampal E-I synapses and found only small effects on average. On one hand, this seems unexpected, since SWRs play an important role in memory consolidation (Joo and Frank, 2018) and the synchronized activity that occurs during SWRs (Buzsáki et al., 1992; Chrobak and Buzsáki, 1996; Nokia et al., 2020; Noguchi et al., 2022) closely resembles LTP protocols (Buzsáki et al., 1987). Moreover, transmission at E-I synapses, in particular, appears to be necessary for SWR generation (Stark et al., 2014). The lack of large effects could potentially be attributed to the fact that the data analyzed here are from animals that are awake, head-fixed and passively viewing visual stimuli. Previous studies on SWRs have involved spatial learning tasks (Girardeau et al., 2009; Ego-Stengel and Wilson, 2010), and often focus on the special role of SWRs during sleep (Chen et al., 2012; Gan et al., 2017). The data here may simply reflect a kind of equilibrium where awake SWRs drive only small changes in efficacy, rather than a larger reorganization that might occur during learning or memory consolidation.

Overall, our results demonstrate that activity-dependent long-term plasticity can partially predict the slow, uncontrolled fluctuations in the efficacy of putative hippocampal E-I synapses measured in large-scale spike recordings. Future studies that examine long-term synaptic changes during learning or memory consolidation in different brain areas may tell us more about the dynamics in synaptic transmission *in vivo*.

## Methods

Here we use the Visual Coding – Neuropixels dataset from the Allen Institute (Dataset: Allen Institute MindScope Program (2019). Allen Brain Observatory -- Neuropixels Visual Coding [dataset] (Siegle et al., 2021). Available from brain-map.org/explore/circuits). During the recordings, head-fixed mice viewed a standardized set of visual stimuli while they were free to run on a wheel. Each recording lasts approximately 2.5 hours and contains 300-900 well-isolated single units from 5 or 6 Neuropixels probes from mouse visual cortex, thalamus, and hippocampus. Here we focus on the putative hippocampal connections within and between areas CA1, CA3, dentate gyrus (DG), subiculum (SUB), and prosubiculum (ProS).

Code for reproducing the methods here is available at https://github.com/ihstevenson/syn_tracking_hc_ei.

### Detection of putative excitatory synaptic connections

We detect putative synaptic connections using the cross-correlograms (CCG) between pairs of neurons

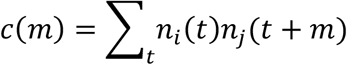

where *n*_*i*_ and *n*_*j*_ denote spike counts for the putative presynaptic *i* and postsynaptic *j* neurons, t denotes time with 0.5ms resolution, and *m* denotes a time interval −20*ms* ≤ *m* ≤ 20*ms*. Given the CCG, we detect putative excitatory synapses in two stages following (Stevenson, 2023). First, we perform an initial screening of neuron pairs that have fast, transient changes in their CCGs. We use a hypothesis test to see whether there are two neighboring bins in the CCG that are statistically significantly different from a convolutional null model of the CCG where fast changes are removed (with *α*=0.1 FDR, using Benjamini– Hochberg linear step-up procedure). Second, we use a model-based approach to fit and screen for putative synaptic effects at specific latencies and timescales. Here, for each pair of neurons that satisfies the initial hypothesis testing criteria we fit an extended GLM based on the model used in (Ren et al., 2020; Stevenson, 2023). Briefly, the modeled spike rate λ at interval *m* in a CCG is given by

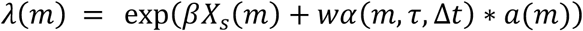

where *X*_*s*_ is a set of cubic B-spline basis functions with equally spaced knots that is weighted by coefficients *β* to describe slow baseline variation in the CCG. *wα*(*m*, τ, Δt) represents the fast, transient synaptic effect, where *w* is the synaptic strength and *α*(*m*, τ, Δt) is an alpha function with a time constant τ and a latency Δt that is then convolved with the auto-correlogram *a*(*m*) of the presynaptic neuron.

We fit the parameters using a penalized maximum likelihood method (Stevenson, 2023) with a Binomial likelihood, and select the neuron pairs that are well described by this parametric model (pseudo-R^2^>0.7) where the CCG has a strong sharp peak (*w* > 0.3, τ < 0.8 *ms*) with a short latency (Δt < 10 ms) and a relatively stable baseline (coefficient of variation of the slow baseline < 0.2).

We generate a model-based estimate of the efficacy and contribution (Levick et al., 1972) using the excess spike probability *eff* = ∑_*m*_Oλ(*m*) – λ_*s*_ (*m*)P where λ_*s*_ denotes the baseline-only model exp(*βX*_*s*_). The standard error for efficacy estimates *se*(*eff*) is estimated by sampling λ(*m*) and λ_*s*_(*m*) via {*β, w*} according to the maximum likelihood estimate and the estimated parameter covariance matrix.

### Estimating time-varying synaptic efficacy

To estimate time-varying efficacy, we first fit the overall parameters for the synaptic effect {*w*, τ, Δt}, then re-estimate *w*_*t*_ based on a set of local spike observations *c*_*t*_ (*m*) generated within a window t − *T* ≤ t ≤ t + *T*.

We optimize the window size *T* by local 5-fold cross-validation. Namely, we estimate *w*_*t*_ based on training observations *c*_*t*_ (*m*) generated using spikes randomly sampled from *n*_(_(*t*)during the window period t −*T* ≤ t ≤ t + *T*. We then evaluate the likelihood (or deviance as in Fig 3B), based on the test set. Here test sets are generated using non-overlapping segments and a fixed 10s window, and predictions are generated from training data with overlapping segments of size *T* with 10s stride. This allows direct comparison of performance across multiple values of *T*. Generally, if the window size *T* is too large, *w*_*t*_ will not accurately follow genuine changes in efficacy, while, if the window size is too small, *w*_*t*_ will track measurement noise.

### Estimating and fitting efficacy change maps

While the model-based coefficients *w*_*t*_ described above provide optimized time-varying estimate of the efficacy fluctuations *eff*(t), we also wish to identify associations between specific environmental events and changes in efficacy Δ*eff*. Here we wish to condition the efficacy estimates based on when the spike observations occur relative to these environmental events. However, since the timescale of the environmental effects is not entirely clear, here we apply a continuous, positive “weight” *r*′ to each presynaptic spike

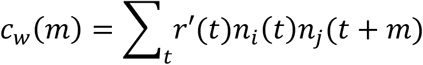

Here we generate *r*(*t*)by filtering the times of the environmental effects with an alpha function

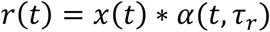

to describe data “after” events at times *x*(*t*)or backward filtering *x*(*t*)* *α*(−t, τ_!_) to describe data “before” events at times *x*(t). Note that τ_*r*_ here is generally much slower than the synaptic time constant τ used above. We additionally normalize *r*, (*t*)= *r*(t)/〈*r*(*t*)〉such that the total number of counts in the weighted CCG can be directly compared to the total for the unweighted CCG. This weighted CCG generalizes previous analyses that have conditioned efficacy estimates based on presynaptic absolute timing (as above) or other features such as the presynaptic inter-spike interval (Swadlow and Gusev, 2001; English et al., 2017; Ghanbari et al., 2020). Here, by fitting the λ(*m*) model to the weighted CCGs, we obtain efficacy estimates before and after specific environmental events.

To describe efficacy changes before and after spike “pairing” events, we first estimate firing rates for the pre and postsynaptic neurons. Here we use bandwidth optimized rate estimates using the methods described in (Shimazaki and Shinomoto, 2010). We then identify percentile ranks for both the pre- and postsynaptic rates and split the rate data into deciles. This generates a joint-rate distribution, along with timings *x*_*pi,pj*_ *(t)*for each combination of pre- and post-synaptic rate deciles *p*_*i*_ and *p*_*j*_ ∈ {1 … 10}. We then estimate an efficacy change map Δ*eff*(*r*_*pre*_, *r*_*post*_) by fitting the corresponding weighted CCG for each pre-post rate combination.

To fit the efficacy change maps, for an initial analysis, we consider learning rules *F*(*r*_*pre*_, *r*_*post*_), and fit the observed Δ*eff*(*r*_*pre*_, *r*_*post*_) using ordinary least squares. We consider a non-Hebbian, linear rule

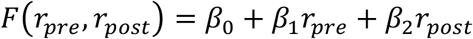

a covariance rule

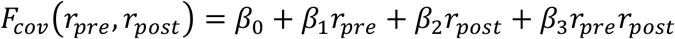

and a BCM rule

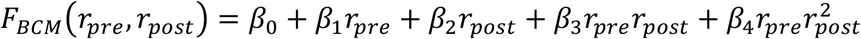

These learning rules differ slightly from their traditional forms. Typically, learning rules assume that *β*_0_ = 0 for stability and that fluctuations in *r* should be represented relative to their means or thresholds. Here we include low order terms to allow greater flexibility in points of reference, and, since the recording length is relatively short there may be some slow drift in efficacy.

### Fitting time-varying efficacy with linear models

Although the efficacy change maps summarize some efficacy changes, it is difficult to determine from these maps alone how different pre-post combinations might interact and difficult to attribute time-varying efficacy fluctuations to unique rate features.

To further determine how efficacy fluctuations are related to the coactivity of the pre- and postsynaptic neurons, we thus use a linear regression fit directly to the estimated efficacy fluctuations *eff*(t). In addition to log-rate covariates (Ren et al., 2022), we add higher order terms to the linear regression to account for accumulated changes that are consistent with the BCM rule

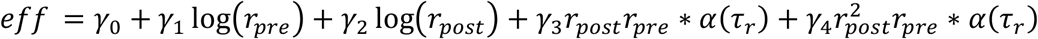

Where the predictors and response variable are all functions of time. Here we use non-overlapping observation windows and optimize *γ* using weighted maximum likelihood estimation where the weights are based on the precision of the time-varying efficacy estimates.

### Sharp-wave ripple detection

To characterize the role of sharp-wave ripples (SWRs) for fluctuations in synaptic efficacy, for each putative synaptic connection, we detect SWRs from the hippocampal local field potential (LFP) from the channel nearest the presynaptic unit. To detect SWRs, we first zero-phase filter the raw LFP signals using a 20th-order bandpass FIR filter (100Hz-250Hz) and Hilbert transform to estimate the envelope power. SWRs are then detected based on the mean and standard deviation (SD) of the peak envelope power (PEP). We classified events that exceed 2.5 SDs and last for more than 15 ms as ripple events. Adjacent events (interval < 15 ms) are merged and expanded. PEP <2 SDs define the start times and end times.

## Acknowledgements

This material is based upon work supported by the National Science Foundation under Grant 1651396. Thanks to the Allen Institute for Brain Science for sharing their datasets and for supporting open science. Thanks to Maxim Volguhsev for helpful discussions.

